# New logic facilitated identification of novel phage recombinase function units

**DOI:** 10.1101/598011

**Authors:** Yizhao Chang, Qian Wang, Tianyuan Su, Qingsheng Qi

## Abstract

Phage recombinase function unit (PRFU) contains one single strand annealing protein (SSAP) and one 5’-3’ exonuclease (EXO). Well-known PRFU such as lambda-Red or Rac RecET has been proved to be useful tool in the recombineering of *Escherichia coli* but had limited efficiency in distant species. Therefore, PRFUs from closely related species were often searched and developed as specific genetic tools. However, there are still many bacteria with few PRFU identified. Here, we hypothesised that the occurrence of PRFU might be related to specific strains. Based on this assumption, we managed to identify 59 unique sets of PRFUs in over 23 species of the genus *Corynebacterium*, a taxonomic group that were reported to be lacking in such system. The identified PRFUs were then classified to show their similarities and relation to species based on both sequential and structural information. Representative PRFUs were verified to be highly effective in mediating recombineering using ssDNA or dsDNA substrates. Analysis of the functional PRFUs indicated that they share similar local genome contexts, which might suggest their common origin. Our findings reveal the relation of PRFUs, species and strains and provide novel candidate genetic manipulation tools in genus *Corynebacterium*.

**Author Summary:** Genus *Corynebacterium* is a taxonomic group with highly diversified species that are of medical, veterinary, or biotechnological relevance. However, being lack of genetic tools limits the study of these species. Our new verified phage recombinase function units (PRFUs) in this genus that are highly effective in mediating recombineering will provide new candidate genetic tools in this genus. Besides, reference genomes were often used for genome study and data mining. While for the mobile genetic elements, their transferable characteristic may affect their occurrence among genomes. Here we provide evidence for this idea by showing that the occurrence of PRFU is strain related but not species related. This may be helpful for the functional genomics study of mobile genetic elements among genomes.

## Introduction

Phage recombinase function unit (PRFU) has been proved to be powerful genetic tool in the recombineering of some bacteria [1–4]. This system mainly contains one single strand annealing protein (SSAP) and one 5’-3’ exonuclease (EXO) [5]. Expression of the SSAP alone can mediate recombineering with single-strand DNA (ssDNA) substrates, while expression of the whole system could use double-strand DNA (dsDNA) as substrates for genome fragment editing[1, 2].

Well-known PRFU such as lambda-Red or Rac RecET has been used for successful recombineering in *Escherichia coli*. The Red system is phage lambda derived, while the RecET system is from the defective prophage Rac of *Escherichia coli* [1]. Lambda-Red contains three adjacent genes gam, bet and exo, which encode a host nuclease inhibiting protein Gam, a SSAP Beta and an EXO accordingly [6]. In *E. coli*, it is possible to mediate recombineering using dsDNA fragment with homology arm as short as 35 base pairs through the expression of Red-EXO, Red-Beta and Red-Gam [7]. RecET, on the other hand, contains only one SSAP RecT and one exonuclease RecE, which has been shown to be highly efficient in linear-linear homologous recombination than lambda-Red [2, 8]. Both lambda-Red and Rac RecET function by specifically interacting with their respective partners, except for the Red-Gam, which is not necessary for dsDNA recombination but can improve the efficiency [1, 9].

Previous studies have showed that PRFU functioned well in related species but performed poorly in distant species, presumably because the exonuclease is more species-specific and could not function well heterogeneously [1, 6]. Thus, the PRFUs of species within the same genus were often searched and developed for specific genetic tool [1, 4, 6, 7]. Typically, analogies of Beta or RecT were often searched, and EXOs besides these SSAPs were also compared [10, 11]. However, several species have been reported to have no such system in this way, such as those in genus *Corynebacterium* [12, 13].

Genus *Corynebacterium* is a taxonomic group with highly diversified species that are of medical, veterinary, or biotechnological relevance[14]. The medical and industrial value encourage the development of genetic manipulation tools for the species of this taxon [13, 15–17]. Up to now, only four species in this genus have been reported to contain PRFU [13, 18]. Even in the well-studied *C. glutamicum*, no PRFU has been identified yet [13].

Here, we proved the idea that the occurrence of PRFU might be related to specific strain by identifying the PRFUs in genus *Corynebacterium*. The identified PRFUs were classified to reveal their relation to species. Typical PRFUs were experimentally verified in mediating recombineering using ssDNA or linear dsDNA substrates. Functional PRFUs were found to share conserved local genome contexts, which might suggest their common origin. These findings provide novel candidate genetic manipulation tools in genus *Corynebacterium* as well as new insight for functional genomics study.

## Results

### Identification of novel PRFU using a database to database method

For most of the studies relating to PRFU, reference genomes were used for bioinformatics analysis [1, 12, 18, 19]. However, as previous verified PRFUs are mainly from prophages or defective prophages, we believe their occurrence may be related to specific bacterial strains. So, all the bacteria genomes as well as the phage genomes of genus *Corynebacterium* in the NCBI genome database were used for study.

As PRFU is mainly composed of one SSAP and one EXO, which usually appear next to each other, we studied the existence of both among genomes to improve prediction accuracy. A database to database method was developed for the searching process. The conserved domain database of SSAPs and EXOs (SSAPdb and EXOdb) were first created using sequences from NCBI conserved domain database. The protein sequences of each genome were then used as queries for PSI-BLAST in the SSAPdb and EXOdb. Then consensus sequences were got through multiple sequence alignment after adding candidate SSAPs or EXOs found to these databases. Then new searches were done until no new SSAP or EXO were found (Fig 1). After iterating for 3 rounds, we identified PRFUs in 84 of the 429 genomes studied, including both classified species and unclassified genomes (S1a-b Fig). As a result, the SSAPs were found in 84 genomes and EXOs were found in 81 genomes. Among them, 54 of the 59 unique SSAPs and 56 of the 57 unique EXOs that were previously annotated as hypothetical protein were identified.

**Fig 1.**
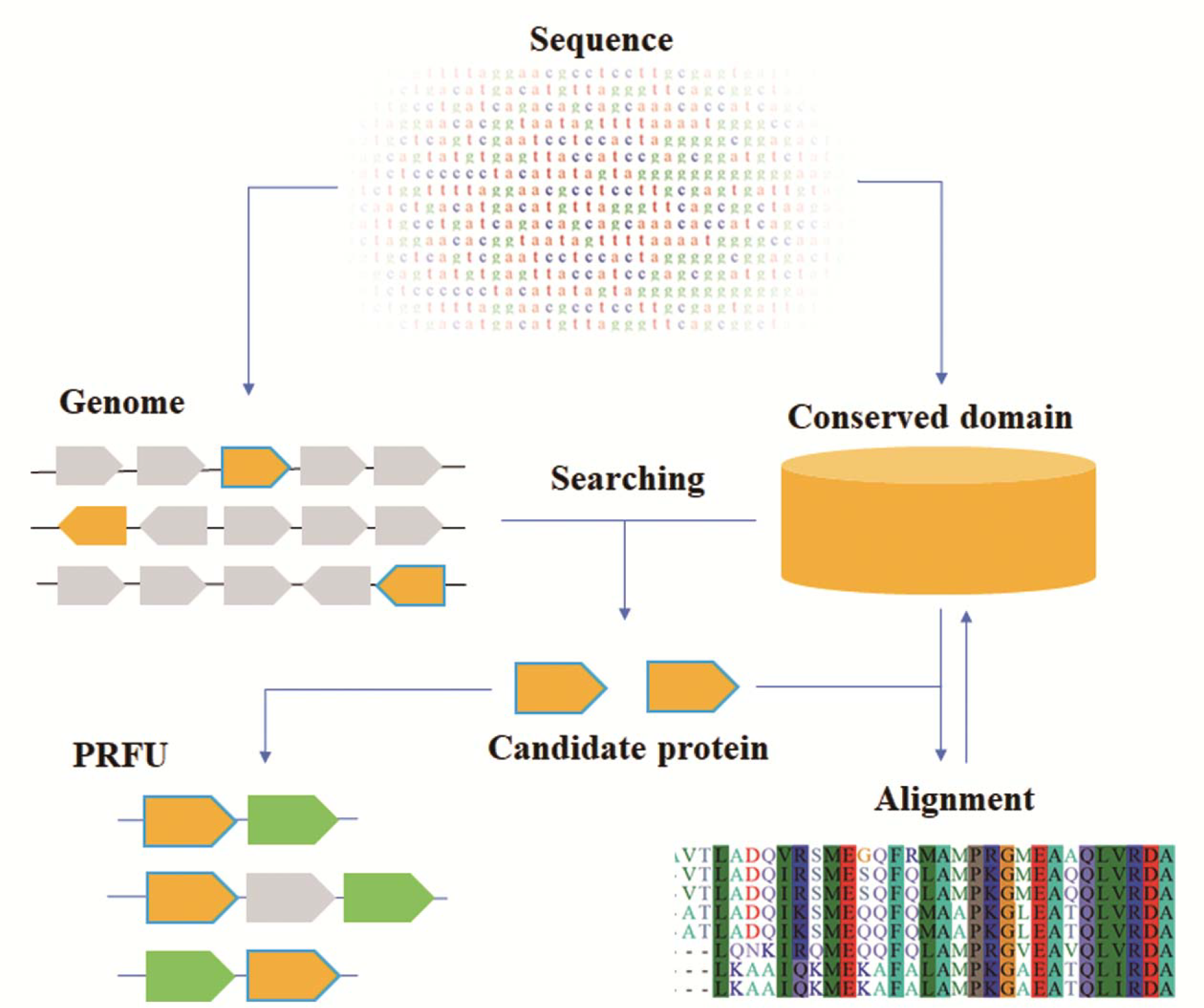
The workflow of finding phage recombinase function unit (PRFU). The genome protein sequence as well as the conserved domain database sequence were first downloaded from NCBI (https://www.ncbi.nlm.nih.gov/). The genome protein sequence database were then searched in the conserved domain database using PSI-BLAST method. Candidate SSAPs or EXOs were then aligned in the conserved domain database to get the consensus parts and form the new conserved domain database. The searching and alignment step were iterated until no new candidates were found. All the candidate SSAPs and EXOs were then collected and paired as PRFUs.

When searching for PRFU among all the plasmids of genus *Corynebacterium*, only one SSAP was found on the plasmid of *C. glutamicum* R strain. This may explain why no PRFU was found previously in *C. glutamicum* when searching its reference genome [12, 13]. In addition to the bacteria genomes, the verified phage genomes of *genus Corynebacterium* were also analyzed but no PRFU was found.

To prove the origin of these PRFUs, the prophages were predicted using an integrated search and annotation tool PHASTER [20]. It was found that among the 84 genomes containing PRFUs, 52 of them have PRFU in the prophage region, while 27 of them have similar proteins coded next to PRFU as those 52 genomes (S1c Fig).

Analysis of the genomes showed that these SSAPs and EXOs form 59 sets of unique PRFUs in over 23 species (Table 1). Most of these species were reported to have no PRFU system or only have SSAP previously [12, 13, 21]. Different from other species, *C. argentoratense* and *C. glutamicum* both separately have one set of incomplete PRFU with only SSAP but no EXO. Besides, not all of the genomes sequenced in these 23 species have PRFU. To test whether this is caused by the incomplete genome sequence data or it just reflects the relationship of PRFUs, genomes and species, the assemble level of the genomes were analyzed. The result shows that the existence of PRFU has no difference between the completely assembled genomes and the not completed ones (S1 Table, t test, p= 0.1314), which indicates that the PRFU occurrence is genome (strain) related but not species related.

**Table 1.**
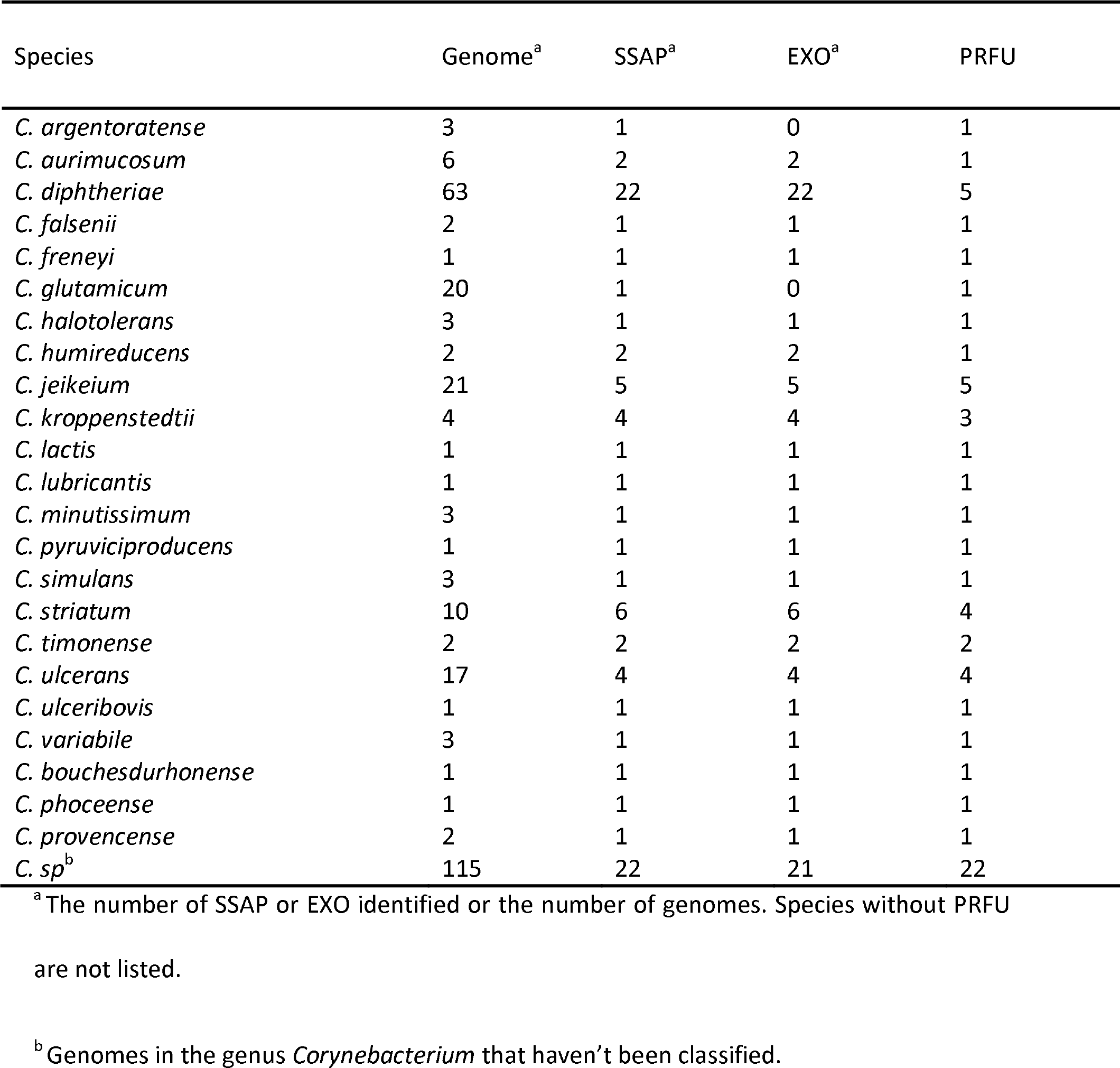
The distribution of PRFUs in genus *Corynebacterium*.

### Classification of the PRFUs and their components

To study the relation of these PRFUs, their component SSAPs and EXOs were classified into 5 types and 3 types separately based on both protein sequence similarity and secondary structure similarity (Figs 2). Each type is conserved in sequence as well as second structure and lies within one branch of the phylogenetic trees. Some identified SSAPs or EXOs from other bacteria were also incorporated.

**Fig 2.**
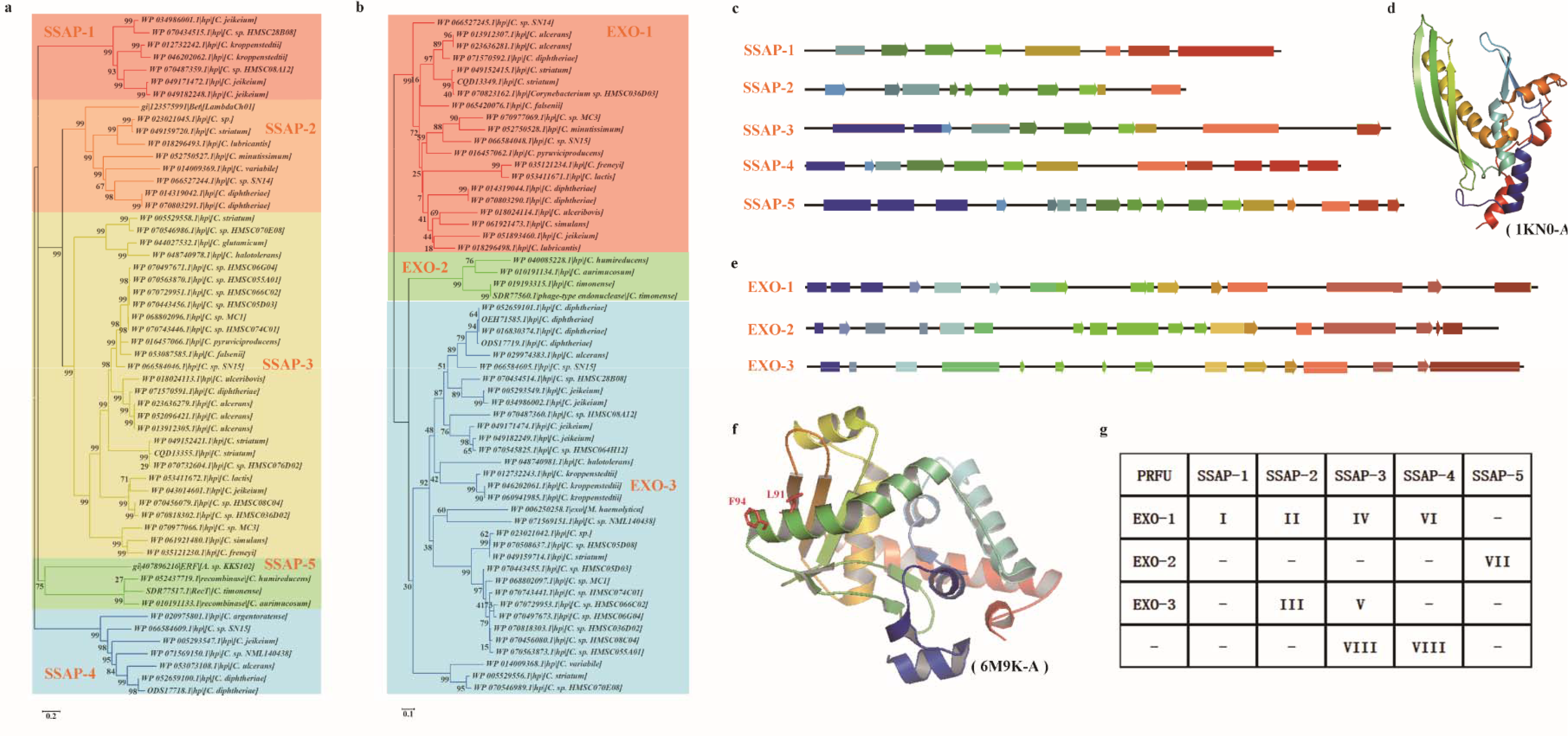
Classification of the PRFUs. **a** Phylogenetic tree and classification of the SSAPs identified. **b** Phylogenetic tree and classification of the EXOs identified. Each type is shown in a different color. The labels are shown as the accession number of the protein, the note for the protein and the source of the protein. The phylogenetic trees were constructed by MEGA6 using Neighbor-Joining method. **c** The schematic second structure of each SSAP types. **d** The N-terminal structure of human SSAP Rad52. PDB: 1KN0. **e** The schematic second structure of each EXO types. f. The structure of λ Exo. PDB: 6M9K. g. The theoretical PRFU type and observed PRFU type. -: not detected.

For SSAPs, the residues in helices are conserved in the same type, while the residues in strands tend to vary from sequence to sequence. Multiple sequence alignment result showed that SSAP-2 shares much similar to Beta, whereas SSAPs in SSAP-5 were found to have sequence similarity with RecT. A conserved β-β-β-α fold structure was observed in the predicted structure of SSAP-1, SSAP-3 and SSAP-4, while a deformed β-β-β-β-β-α fold structure was observed in SSAP-2 and SSAP-5 (Fig 2c). The same β-β-β-α fold has been observed in human Rad52, which was found to be composed of highly conserved amino acid residues that are responsible for ssDNA and dsDNA binding [22]. The structure of the N-terminal of human Rad52 (Rad52_1-212_) was shown in Fig 2d as an indicant. Actually, a deformed β-β-α-β-β-β-α fold of Rad52_1-212_ was observed in all SSAP types except SSAP-1. Besides, similar to the Rad52_1-212_, both the C-terminal and the N-terminal of the SSAPs are rich in helices. However, compared to the conserved N-terminal, the residues in the C-terminal region of different SSAP types are irregular. This might be accordingly with the fact that the conserved N-terminal of SSAP is responsible for homologous pairing, while the C-terminal interacts with EXO or host factors [23–26].

Unlike SSAPs, the second structure of EXOs varied a lot from type to type (Fig 2e). The residues in the N-terminal half of EXOs are conserved across types, while the residues in the C-terminal half vary a lot even in the same type. The residue Leu-91 and residue Phe-94 of λ Exo have been found to be important for forming the λ Exo-Beta complex [26]. These residues lie within a helix of the α-α-β-β fold of the λ Exo (Fig 2f), which was also observed in EXO-3.

These SSAPs and EXOs form eight types of PRFU in total, which is less than their theoretical combination (Fig 2g). Typically, SSAP-1 only appear with EXO-1 and forms PRFU-I, while SSAP-5 and EXO-2 only appear with each other and form PRFU-VII. Interestingly, all the species that harbour more than one type of PRFU are pathogenic (S2 Fig), while not all pathogenic species have multi-type PRFUs.

### Verification of the function of PRFUs in vivo

Previous study indicated that the expression of SSAP alone could promote ssDNA mediated recombineering, while expression of the full PRFU would facilitate genetic manipulation of genome with dsDNA [1, 6, 7, 27]. To verify the function of PRFUs identified, both oligo mediated recombination and dsDNA mediated recombination experiments were conducted in a previous constructed strain C.g-kan(−) [28].

To verify the function of SSAPs, eight SSAPs were expressed on the shuttle expression vector PXMJ19 (Fig 3a). The SSAPs were chosen based on the species evolutional relationship (inferred from the phylogenetic tree of their 16s rRNA, S3 Fig) as well as the PRFU type. An oligo targeting the defective kan(-) was used for ssDNA mediated recombineering to recover the function of kan (Fig 3b, [13, 28]). As a result, five of them could promote ssDNA mediated recombination in *C. glutamicum* (Fig 3c-d, Table 2 Sap1-Sap8). While the expression of WP_010191133.1 from *C. aurimucosum* got the highest recombinant number and the highest recombination frequency, the recombination frequency of WP_052659100.1 from C. diphtheriae and WP_044027532.1 from *C. glutamicum* was similar to that of the positive control RecT from *E. coli*. No or extremely low level of recombination phenomenon was observed in strains expressing WP_060941986.1 from *C. Kroppenstedtii*, WP_018296493.1 from *C. lubricantis*, and WP_049159720.1 from *C. striatum*.

**Table 2.**
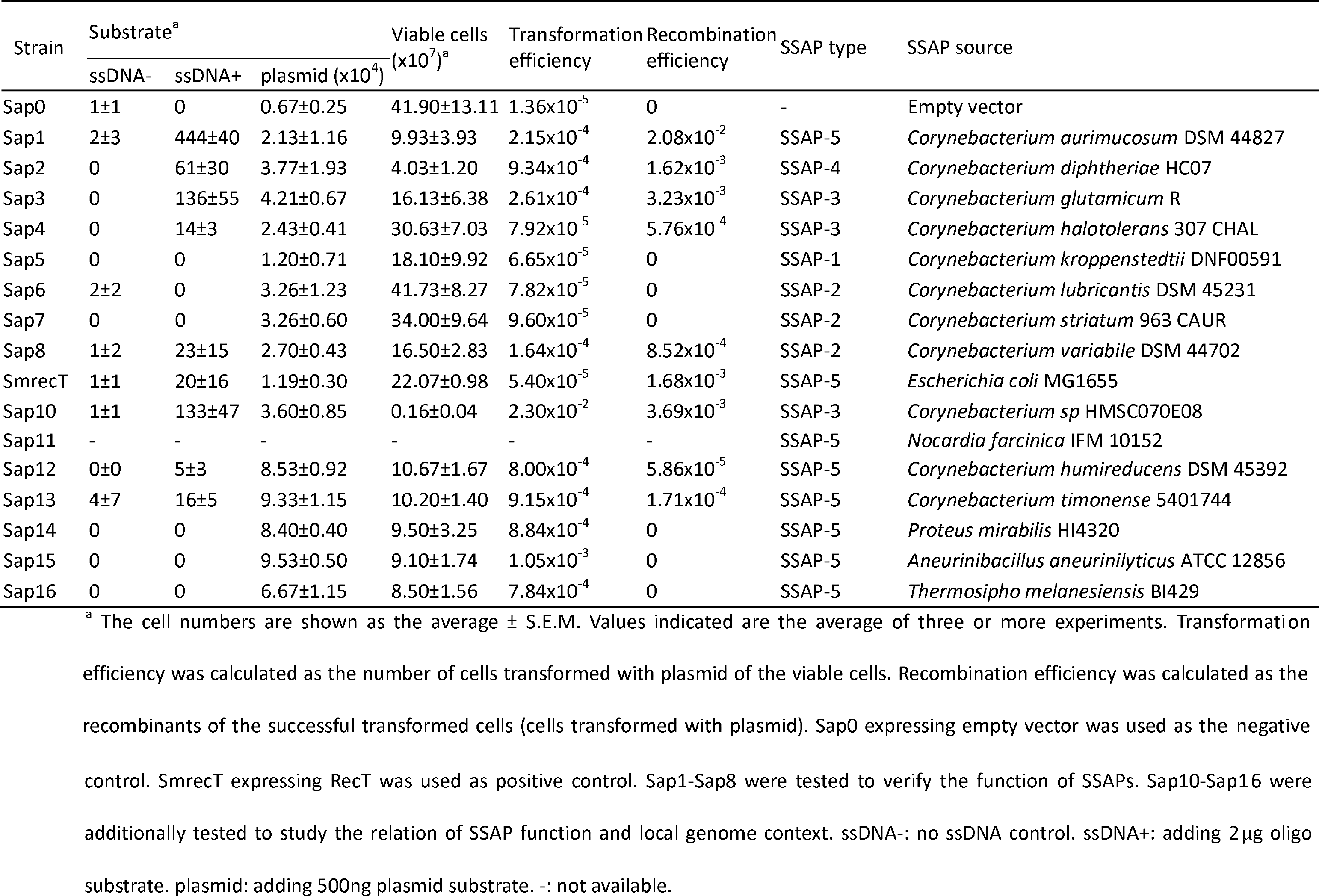
Recombineering with ssDNA using different SSAPs.

**Fig 3.**
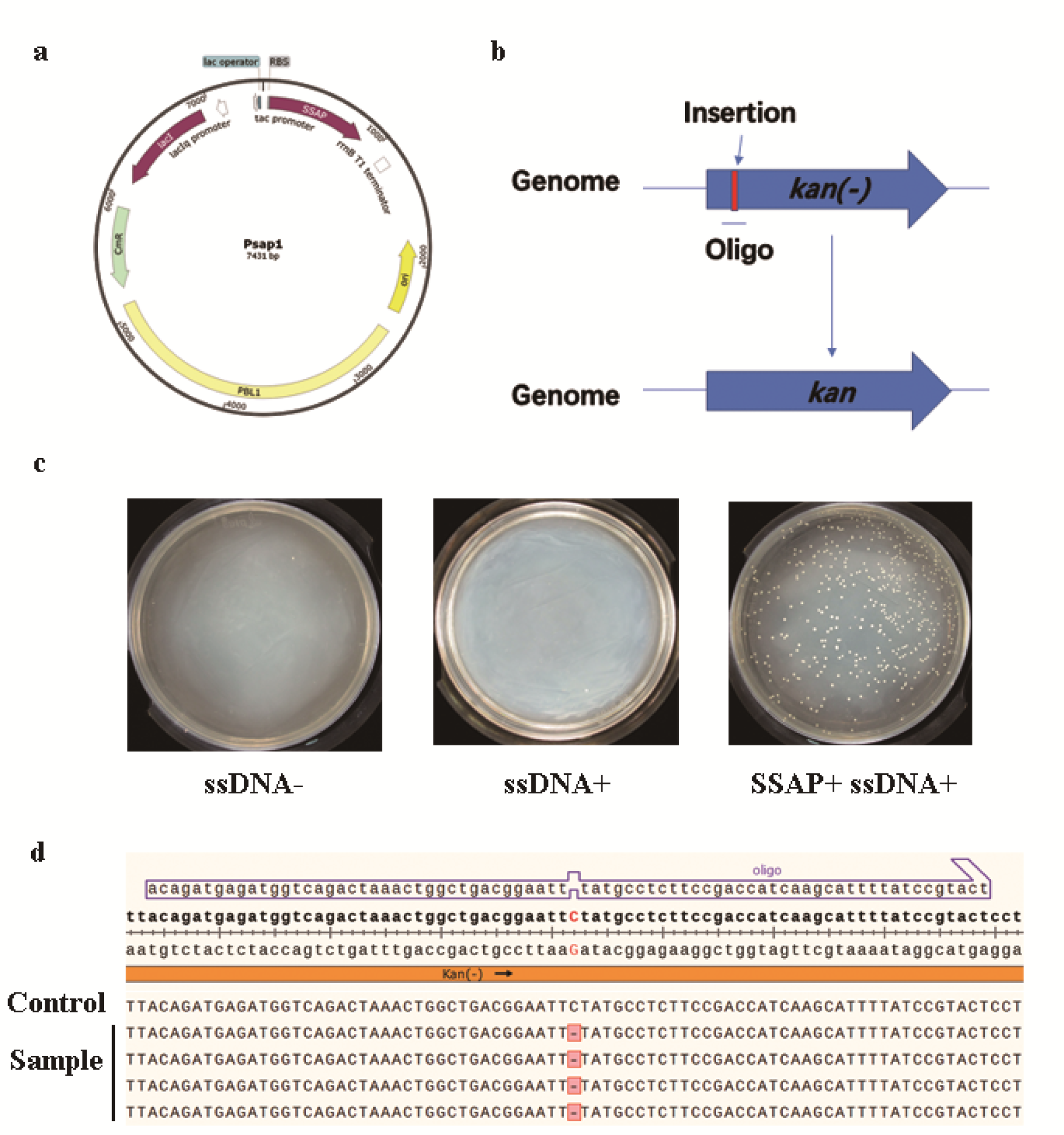
Recombineering using SSAP with oligo. **a** The plasmid used for SSAP expression. **b** Schematic of oligo mediated recombineering. **c** The culture of no ssDNA control (ssDNA-), no SSAP control (SSAP+) and recombinants survival the selection (SSAP+ ssDNA+). **d** Sequence evidence for four recombinants sample and one no recombineering control.

To verify the recombineering ability of the full PRFUs whose SSAPs were proved to be functional above, their component EXOs were expressed after the corresponding SSAPs (Fig 4a). Selection marker interspaced by 800bp homology arm targeting recA was used as dsDNA substrate (Fig 4b). Though no complete PRFU was found in *C. glutamicum* R strain, the gene next to its SSAP coding gene was also tested. It was found that both PRFU from *C. aurimucosum* and RecET could promote highly efficient recombination in vivo with the recombination frequency at 10^−2^ magnitude (Table 3). The PRFU from *C. halotolerans* was also proved to be functional, while was an order of magnitude less efficient. No or extremely low recombination frequency was observed in the PRFUs from *C. glutamicum* and *C. diphtheriae*. The experimental evidences were shown in Fig 4c-e.

**Table 3.** Recombineering with dsDNA using different PRFUs.

| Strain | Substrate<sup>a</sup> |  |  | Viable cells<br>(x10<sup>5</sup>)<sup>a</sup> | Transformation<br>efficiency | Recombination<br>frequency | PRFU<br>type | PRFU source |
| --- | --- | --- | --- | --- | --- | --- | --- | --- |
|  | dsDNA- | dsDNA+ | plasmid<br>(x10<sup>5</sup>) |  |  |  |  |  |
| ET1 | 0 | 2365±822 | 1.52±0.09 | 3.73±1.44 | 4.08x10<sup>-1</sup> | 1.56x10<sup>-2</sup> | VII | *Corynebacterium aurimucosum* DSM 44827 |
| ET2 | 0 | 4±2 | 3.11±2.07 | 6.10±1.87 | 5.1x10<sup>-1</sup> | 1.29x10<sup>-5</sup> | VI | *Corynebacterium diphtheriae* HC07 |
| ET3 | 0 | 0 | 1.53±0.50 | 8.5±1.97 | 1.8x10<sup>-1</sup> | 0 | VIII | *Corynebacterium glutamicum* R |
| ET4 | 0 | 85 ±73 | 0.66±0.23 | 11.43±8.44 | 5.73x10<sup>-1</sup> | 1.3x10<sup>-3</sup> | IV | *Corynebacterium halotolerans* 307 CHAL |
| ET8 | - | - | - | - | - | - | III | *Corynebacterium variabile* DSM 44702 |
| ET9 | 1 | 2456<br>±391 | 1.02±0.17 | 6.03±1.42 | 1.69x10<sup>-1</sup> | 2.41x10<sup>-2</sup> | - | *Escherichia coli* MG1655 |

<sup>a</sup> The cell numbers are shown as the average ± S.E.M. Values indicated are the average of three or more experiments. Transformation efficiency was calculated as the number of cells transformed with plasmid of the viable cells. Recombination efficiency was calculated as the recombinants of the successful transformed cells (cells transformed with plasmid). ET9 expressing RecET was used as the positive control. dsDNA-: no dsDNA control. dsDNA+: adding 1µg dsDNA substrate. plasmid: adding 500ng plasmid substrate. -: not available.

**Fig 4.**
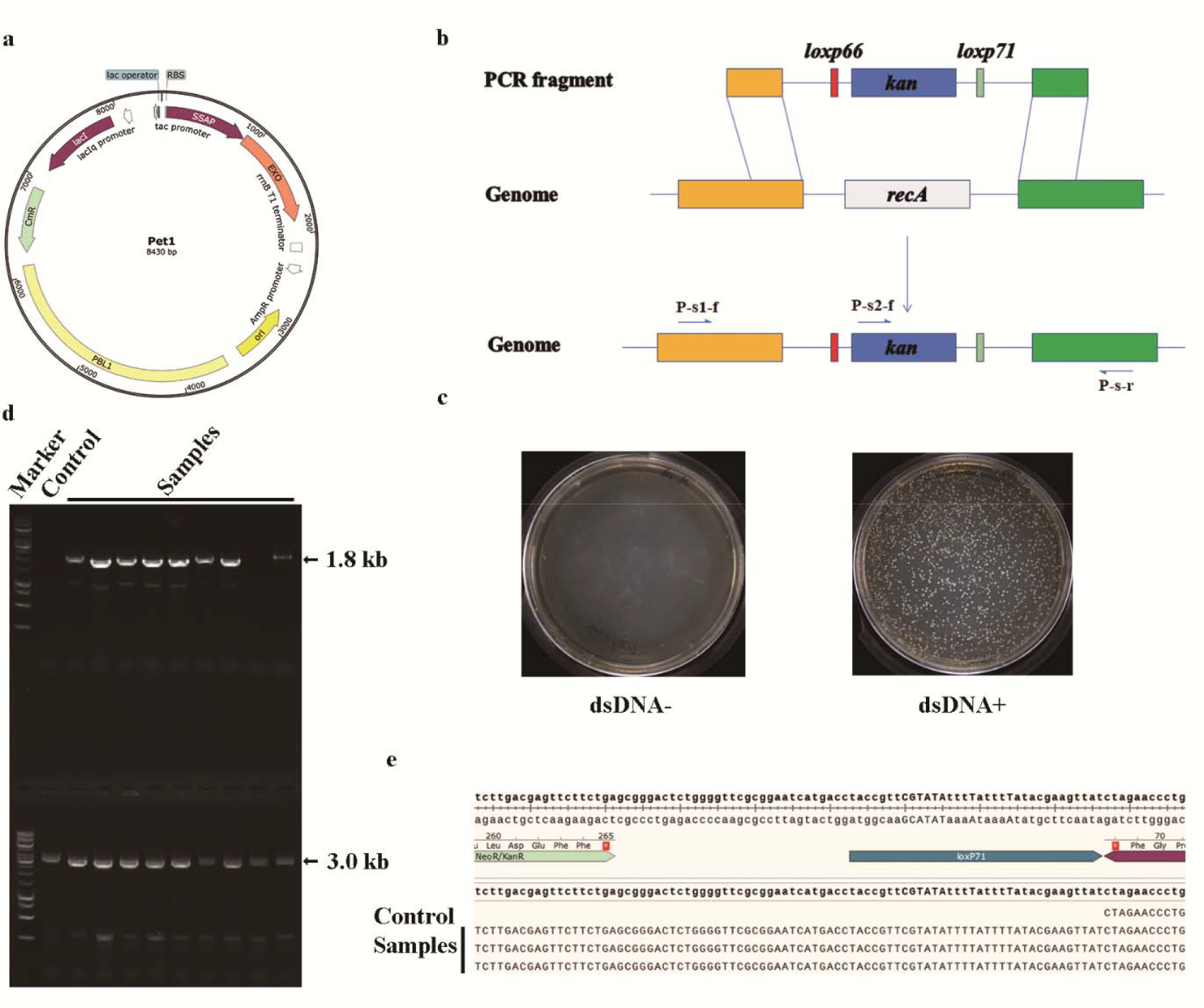
Recombineering using PRFU with dsDNA substrates. **a** The plasmid used for PRFU expression. **b** Schematic of dsDNA mediated recombineering. **c** The culture of no dsDNA control (dsDNA-) and recombinants survival the selection (dsDNA+). **d** Gel electrophoresis evidence of 9 samples and one no recombineering control. The 1.8 kb line was amplified by primers P-s2-f and P-s-r. The 3.0 kb line was amplified by primers P-s1-f and P-s-r. **e** Sequence evidence for three recombinants sample and one no recombineering control.

## Discussion

Although genome sequence grew rapidly these years, few PRFUs have been identified. This and the phage origin of verified PRFUs led us to investigate all the genomes sequenced in genus *Corynebacterium*. As each genome represent a strain record, the occurrence of PRFUs in both reference and non-reference genomes and the irregular distribution pattern of PRFUs among genomes within the same species verified the idea that the existence of PRFUs is strain related but not species related. This indicates that the prediction of PRFU using reference genomes might lead to error conclusion about their existence in the species.

For most of the time, sequence homology-based method was used for the prediction of protein function [29]. When predicting SSAP, a single query such as RecT or Red-Beta was used to start the search for new homologies[13, 19]. To improve the prediction accuracy, the protein sequences of genomes were searched for several rounds in the conserved domain database of SSAPs or EXOs in this study. This enabled us to select results that were identical to several records in the conserved domain database. Besides, by doing iteration, distant-related SSAPs or EXOs were also identified.

To test the function of PRFUs found in this study, we verified several of them with both ssDNA and linear dsDNA mediated recombineering heterogeneously in *C. glutamicum*. Several tested PRFUs were proved to be functional in promoting ssDNA or linear dsDNA mediated recombination. Actually, some PRFUs tested showed relatively high ability in conducting recombination as previously reported tools in other bacteria [1, 6, 7]. As these PRFUs can function in *C. glutamicum*, they are believed to function even better in their hosts for their specific molecular interactions with the host factors [1]. This suggested their great potential for recombineering in genus *Corynebacterium*.

Four of the five functional SSAPs verified in this study have more closed local genome context relation than others and lie within the same branch of the local genome context tree (Fig 5a). All of the S_dis_ of local genome context to *C. glutamicum* R strain in this branch are smaller than 35 and have a wider range from 0 to 35 compared to others, which have a narrow range of S_dis_ from 35 to 40 (Fig 5b). This suggests that the local genome context conservation might be related to the function of SSAPs. Additional verified SSAPs with S_dis_ smaller than 35 functioning in the recombineering experiment verified this idea (Table 2, Fig 5). Analysis of the components of the local genome contexts in this branch indicates that they are rich in genes whose products may interact with DNA, such as transcriptional regulators and site-specific integrase. Besides, no linear relation between the species 16S rRNA distance and local genome context distance was observed (S4 Fig). These suggest that the local genome context conservation in this branch might be caused by the horizontal gene transfer.

**Fig 5.**
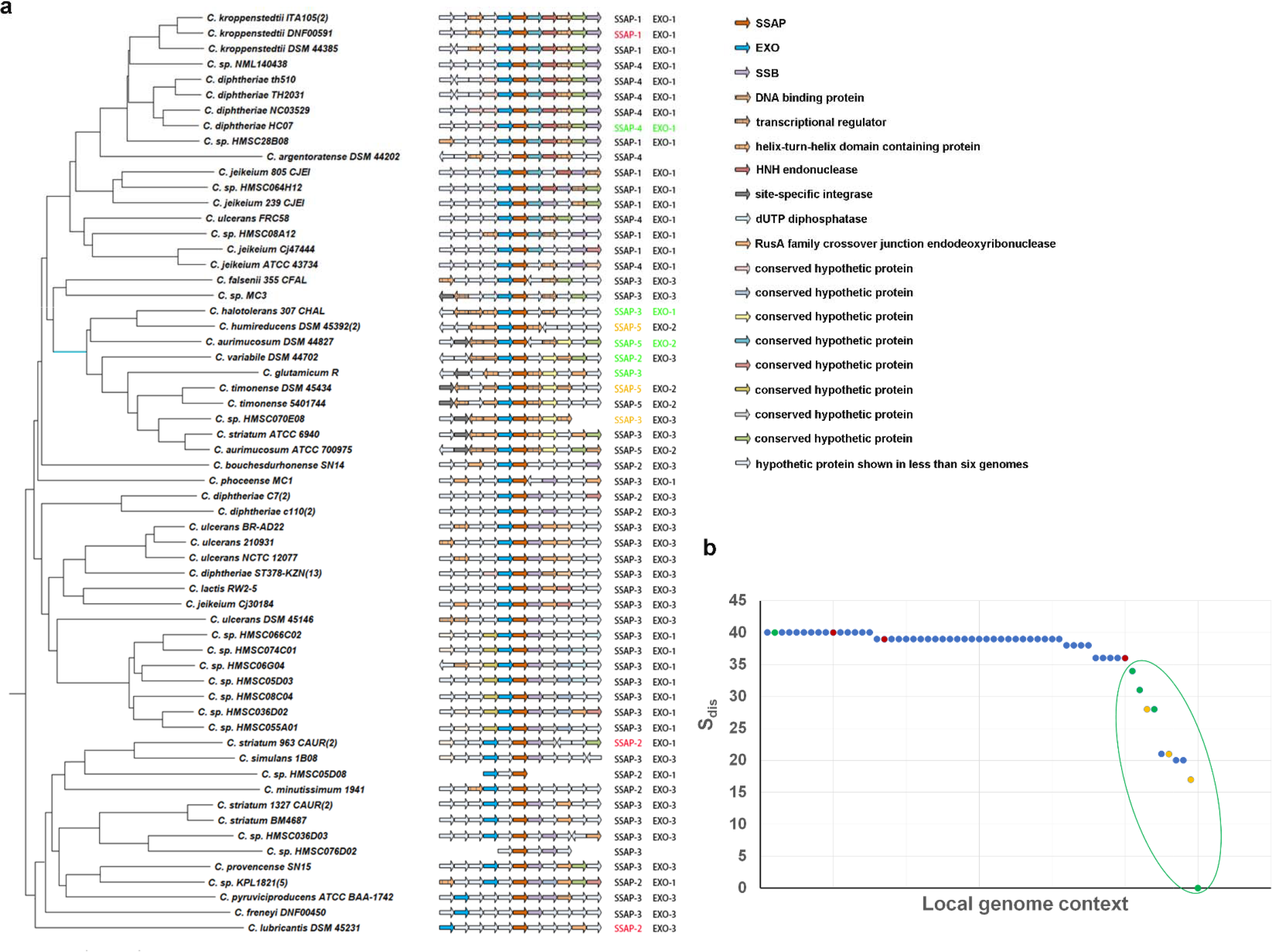
Local genome context of the PRFUs. **a** The phylogenetic tree of local genome context of PRFU. **b** The local genome context distance (S_dis_) of different species with PRFUs to that of *C. glutamicum*. pRFUs functioned in the recombineering experiment were colored in green. Those showed no function were colored in red. Additional verified functional SSAPs were colored in orange.

In this study, the database to database searching method facilitated accurate prediction of novel PRFUs in the species that were reported to have no such system. Analysis of the distribution of PRFUs among genomes indicates that their occurrence is related to strains but not species. This may provide new insights for the study of mobile genetic elements. Besides, several PRFUs were proved to be highly effective in mediating recombineering using ssDNA or dsDNA substrates in this study. Due to the limited genetic manipulation tools available in genus **Corynebacterium**, the PRFUs verified will also provide new candidate tools in these and related species.

## Materials and methods

### Analysis of PRFU across genomes

Genomes of genus *Corynebacterium*, including chromosomes, plasmids and phages were downloaded from NCBI (https://www.ncbi.nlm.nih.gov/) genome database. The conserved domain data of SSAP and EXO from the Conserved Domains Database of NCBI (Accession number: cl04285, cl04500, cl01936 and pfam09588) were also downloaded. Then the following process were iterated:

1. Multiple sequence alignments were done to get the consensus of SSAPs and EXOs and construct database SSAPdb as well as EXOdb.
2. Then genomes of genus *Corynebacterium* were searched for potential SSAP and EXO in the SSAPdb and EXOdb separately through PSI-BLAST for one iteration.
3. Potential SSAP and EXO were extracted and added to the SSAPdb or EXOdb accordingly.

This process was iterated for three times until no new potential SSAP or EXO was found. Potential SSAPs were chosen from the result of PSI-BLAST using a cut off E value of 10^−6^. Potential EXOs were chosen by setting a cut off E value as 10^−5^.

Prophage was predicted by PHASTER (http://phaster.ca/) [20]. The genome location of PRFUs as well as the local genome context were used for the verification of their prophage origin.

### Protein sequence and structure analysis

Multiple protein sequence alignment was performed with MUSCLE [30]. The alignment results were then edited manually to get the consensus parts and shown as the default settings in ClustalX [31].

All of the second structures of the alignments were predicated by Jpred4 (http://www.compbio.dundee.ac.uk/jpred4/index.html) [32].

### Alignment of the local genome context

The adjacent five proteins coded upstream and downstream the SSAP of each genome were extracted to evaluate the local genome context similarity if available. First, the similarity of these proteins was analyzed through PSI-BLAST. Protein pairs with E value less than 10^−5^ were grouped together. The Groups with proteins of similar annotation or the proteins that were not grouped in the previous step but have similar annotation were merged into one group. The proteins that were not grouped yet were each assigned a group separately.

A similarity matrix of these local genome context was then calculated through the similarity score S_sim_ between each pair of them. The similarity score S_sim_ contained two parts, S_sam_ and S_loc_. S_sam_ was calculated as the sum number of protein pairs from the two genomes that are of same group. S_loc_ was the sum of S_i_, which was calculated in the following way: For each protein coded upstream SSAP and downstream SSAP, its position was assigned a site i. Site i has a range from 1 to 11, corresponding to the position from the farthest upstream the position of SSAP to the farthest downstream. If the two proteins of both genome at the site i belongs to the same group, then S_i_ was assigned 1,2,3,4,5,6,5,4,3,2 or 1 for each i from 1 to 11 accordingly. This can be written as:

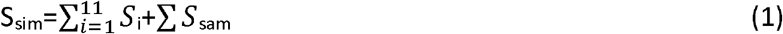

The distance matrix of these genomes was got from the similarity matrix, which used a score S_dis_. S_dis_ was calculated as:

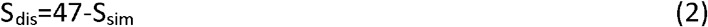

The number 47 was the S_sim_ for a pair of completely same local genome context that had an ideal S_dis_ value of 0. As different site was assigned a different weight, we call these site-weighted scores.

### Phylogenetic analysis

The genome phylogenetic tree was constructed using the 16S rRNA of the representative genomes. The protein phylogenetic tree was constructed using the multiple sequence alignment results of SSAPs and EXOs. For the construction of local genome context phylogenetic tree, a distance matrix was used as input. The phylogenetic trees were constructed by MEGA6 using Neighbor-Joining method with the default settings except for the Interior-branch Test value, which is set as 1000 [33].

### Strain cultivation

The *C.glutamicum* strain C.g-kan(−) was cultured in LB or BHIS medium. For the preparation of the electrocompetent cells, the strains were first cultured in LB medium overnight at 30°C. The cultures were then added to 100ml BHIS medium at a start optical density of 0.3 and cultured at 30 °C until the optical density reached 0.9, with IPTG and antibiotics added according to the circumstance. The culture was then chilled on ice for 30min and centrifuged at 4°C to collect the cells. The sediments were washed for four times using 30ml 10% (v/v) glycerol and resuspended in 1ml 10% (v/v) glycerol. Then a volume of 100ul each tube was divided and stored at −80°C for later experiment. To maintain the plasmids, a final concentration of 100 μg/ml spectinomycin, 15µg/ml kanamycin or 10 μg/ml chloromycetin was added to the corresponding cultures. For induction purpose, a final concentration of 0.25 mmol/L IPTG was used [34].

### Statistics

The ratio of the number of sequence finished genomes with PRFU and sequence finished genomes as well as the ratio of the number of sequence unfinished genomes with PRFU and sequence unfinished genomes were used for t-test (S1 Table).

### Recombineering with PRFUs

The plasmid, ssDNA and dsDNA substrates were transformed into *C. glutamicum* electrocompetent cells by electroporation. Different substrates were pre-chilled and added to the electrocompetent cells according to the circumstances. For recombineering with ssDNA, 2µg oligo was added to the competent cells. For plasmid and dsDNA, 500ng or 1µg substrate was used. The mixture was then used for electroporation at 1.8kv/mm, 200Ω, 25µF. Then 1ml 46°C BHIS medium was instantly added. The mixture was heat shocked at 46 °C for 6min and then cultured for outgrowth at 30 °C, 200rpm up to 1h for strain construction purpose or 30 °C, 150rpm for 5h for the transformation of ssDNA, dsDNA and control plasmid. Cultures were then plated directly after collecting the cells for most of the time. To count the number of viable cells, 0.1ml of 10^−4^ dilution was plated on BHIS with chloromycetin added [13, 35].

Fragments were prepared through overlap extension PCR using TransStart FastPfu Fly DNA Polymerase (TransGen, AP231-03) and collected using DNA Gel Extraction Kit (TsingKe, GE101-200). Results were verified by agarose gel electrophoresis and Sanger sequence.

## Acknowledgements

We thank Haiying Jin for her help in the recombineering with ssDNA experiment. We also thank Qilong Qin for reviewing the manuscript.

## Author contributions

YZ designed and conducted the experiment, prepared and revised the manuscript. QS designed, supervised the study and revised the manuscript. QW revised the manuscript. TY participated the experiment.

## Supporting information

**S1 Fig. The PRFUs in genus *Corynebacterium*. a** The genomes with or without PRFUs. **b** The genomes in species with or without PRFU. **c** The phage origin of predicted PRFUs. F: genomes with PRFU, NF: genomes without PRFU. CL: PRFUs found in genomes that has been classified into species, UCL: PRFUs found in genomes that hasn’t been classified into any species, F: genomes with PRFU, NF: genomes wiithout PRFU. Numbers indicate the genome number.

**S2 Fig. The distribution of different PRFU type among species.** Pathogenetic species were colored in red.

**S3 Fig. The 16S rRNA phylogenetic tree of genus *Corynebacterium*.** The 16S rRNA of the representative genomes of classified species were used to generate the tree. Sequences from *Mycobacteria* were used as outgroup. The phylogenetic trees were constructed by MEGA6 (2) using Neighbor-Joining method with the default settings except for the Interior-branch Test value, which is set as 1000.

**S4 Fig. Colinear analysis of the local genome context distance and species 16S rRNA distance.** PRFUs functioned in the recombineering experiment were colored in green. Those showed no function were colored in red.

**S1 Table. The occurrence of PRFU and genome assemble level**

**S2 Table. Strains and plasmids used in the study**

