## Supplementary material for "New logic facilitated identification of novel phage recombinase function units": S1 Table

| **Table S1. The occurrence of PRFU and genome assemble level** | | | | | | | | | |
| --- | --- | --- | --- | --- | --- | --- | --- | --- | --- |
| Species | Genome^a^ | SSAP^a^ | EXO^a^ | Finished genome with PRFU^a^ | Finished genome^a^ | F_ratio^c^ | Unfinished genome with PRFU^a^ | Unfinished genome^a^ | NF_ratio^d^ |
| *C. argentoratense* | 3 | 1 | 0 | 1 | 1 | 1.00 | 0 | 2 | 0.00 |
| *C. aurimucosum* | 6 | 2 | 2 | 1 | 1 | 1.00 | 1 | 6 | 0.17 |
| *C. sp^b^* | 119 | 21 | 20 | 0 | 2 | 0.00 | 25 | 117 | 0.21 |
| *C. diphtheriae* | 63 | 22 | 22 | 1 | 14 | 0.07 | 21 | 49 | 0.43 |
| *C. falsenii* | 2 | 1 | 1 | 0 | 1 | 0.00 | 1 | 1 | 1.00 |
| *C. freneyi* | 1 | 1 | 1 | 0 | 0 | - | 1 | 1 | 1.00 |
| *C. glutamicum* | 20 | 1 | 0 | 1 | 15 | 0.07 | 0 | 5 | 0.00 |
| *C. halotolerans* | 3 | 1 | 1 | 0 | 1 | 0.00 | 1 | 2 | 0.50 |
| *C. humireducens* | 2 | 2 | 2 | 1 | 1 | 1.00 | 1 | 1 | 1.00 |
| *C. jeikeium* | 21 | 5 | 5 | 0 | 1 | 0.00 | 5 | 20 | 0.25 |
| *C. kroppenstedtii* | 4 | 4 | 4 | 1 | 1 | 1.00 | 3 | 3 | 1.00 |
| *C. lactis* | 1 | 1 | 1 | 1 | 1 | 1.00 | 0 | 0 | - |
| *C. lubricantis* | 1 | 1 | 1 | 0 | 0 | - | 1 | 1 | 1.00 |
| *C. minutissimum* | 3 | 1 | 1 | 0 | 0 | - | 1 | 3 | 0.33 |
| *C. pyruviciproducens* | 1 | 1 | 1 | 0 | 0 | - | 1 | 1 | 1.00 |
| *C. simulans* | 3 | 1 | 1 | 0 | 2 | 0.00 | 1 | 1 | 1.00 |
| *C. striatum* | 10 | 6 | 6 | 0 | 0 | - | 6 | 10 | 0.60 |
| *C. timonense* | 2 | 2 | 2 | 0 | 0 | - | 2 | 2 | 1.00 |
| *C. ulcerans* | 17 | 4 | 4 | 3 | 10 | 0.30 | 1 | 7 | 0.14 |
| *C. ulceribovis* | 1 | 1 | 1 | 0 | 0 | - | 1 | 1 | 1.00 |
| *C. variabile* | 3 | 1 | 1 | 1 | 1 | 1.00 | 0 | 2 | 0.00 |
| *C. bouchesdurhonense* | 1 | 1 | 1 | 0 | 0 | - | 1 | 1 | 1.00 |
| *C. phoceense MC1* | 1 | 1 | 1 | 0 | 0 | - | 1 | 1 | 1.00 |
| *C. provencense* | 2 | 2 | 2 | 1 | 1 | 1.00 | 1 | 1 | 1.00 |
| ^a^ The number genomes. | | | | | | | | | |
| ^b^ Genomes in genus *Corynebacterium* that has not been classified into species. | | | | | | | | | |
| ^c^ The ratio of the number of finished genome with PRFU and finished genomes. | | | | | | | | | |
| ^d^ The ratio of the number of unfinished genome with PRFU and unfinished genomes. | | | | | | | | | |
