## Supplementary material for "New logic facilitated identification of novel phage recombinase function units": S2 Table

| **Table S2. Strains and plasmids used in the study** | | |
| --- | --- | --- |
| Name | Note | Source |
| C.g-kan(−) | *Corynebacterium glutamicum* 13032, *kan(-)* | lab stock |
| Sap0 | Cglnk expressing PXMJ19 | this study |
| Sap1 | Cglnk expressing Psap1 | this study |
| Sap2 | Cglnk expressing Psap2 | this study |
| Sap3 | Cglnk expressing Psap3 | this study |
| Sap4 | Cglnk expressing Psap4 | this study |
| Sap5 | Cglnk expressing Psap5 | this study |
| Sap6 | Cglnk expressing Psap6 | this study |
| Sap7 | Cglnk expressing Psap7 | this study |
| Sap8 | Cglnk expressing Psap8 | this study |
| Sap9 | Cglnk expressing Psap9 | this study |
| Sap10 | Cglnk expressing Psap10 | this study |
| Sap11 | Cglnk expressing Psap11 | this study |
| Sap12 | Cglnk expressing Psap12 | this study |
| Sap13 | Cglnk expressing Psap13 | this study |
| Sap14 | Cglnk expressing Psap14 | this study |
| Sap15 | Cglnk expressing Psap15 | this study |
| Sap16 | Cglnk expressing Psap16 | this study |
| ET1 | Cglnk expressing ET1 | this study |
| ET2 | Cglnk expressing ET2 | this study |
| ET3 | Cglnk expressing ET3 | this study |
| ET4 | Cglnk expressing ET4 | this study |
| ET8 | Cglnk expressing ET5 | this study |
| ET9 | Cglnk expressing ET6 | this study |
| ET10 | Cglnk expressing ET7 | this study |
| ET12 | Cglnk expressing ET8 | this study |
| ET13 | Cglnk expressing ET9 | this study |
| PXMJ19 | *C. glutamicum*-*E. coli* shuttle expression vector,CmR | lab stock |
| pECXK99E | *C. glutamicum*-*E. coli* shuttle expression vector,KanR | lab stock |
| Psap1 | PXMJ19 expressing SSAP from *Corynebacterium aurimucosum* DSM 44827 | this study |
| Psap2 | PXMJ19 expressing SSAP from *Corynebacterium diphtheriae* HC07 | this study |
| Psap3 | PXMJ19 expressing SSAP from *Corynebacterium glutamicum* R | this study |
| Psap4 | PXMJ19 expressing SSAP from *Corynebacterium halotolerans* 307 CHAL | this study |
| Psap5 | PXMJ19 expressing SSAP from *Corynebacterium kroppenstedtii* DNF00591 | this study |
| Psap6 | PXMJ19 expressing SSAP from *Corynebacterium lubricantis* DSM 45231 | this study |
| Psap7 | PXMJ19 expressing SSAP from *Corynebacterium striatum* 963 CAUR | this study |
| Psap8 | PXMJ19 expressing SSAP from *Corynebacterium variabile* DSM 44702 | this study |
| Psap9 | PXMJ19 expressing SSAP from *Escherichia coli* MG1655 | this study |
| Psap10 | PXMJ19 expressing SSAP from *Corynebacterium sp* HMSC070E08 | this study |
| Psap11 | PXMJ19 expressing SSAP from *Nocardia farcinica* IFM 10152 | this study |
| Psap12 | PXMJ19 expressing SSAP from *Corynebacterium humireducens* DSM 45392 | this study |
| Psap13 | PXMJ19 expressing SSAP from *Corynebacterium timonense* 5401744 | this study |
| Psap14 | PXMJ19 expressing SSAP from *Proteus mirabilis* HI4320 | this study |
| Psap15 | PXMJ19 expressing SSAP from *Aneurinibacillus aneurinilyticus* ATCC 12856 | this study |
| Psap16 | PXMJ19 expressing SSAP from *Thermosipho melanesiensis* BI429 | this study |
| Pet1 | PXMJ19 expressing SSAP from *Corynebacterium aurimucosum DSM 44827* | this study |
| Pet2 | PXMJ19 expressing SSAP from *Corynebacterium diphtheriae HC07* | this study |
| Pet3 | PXMJ19 expressing SSAP from *Corynebacterium glutamicum R* | this study |
| Pet4 | PXMJ19 expressing SSAP from *Corynebacterium halotolerans 307 CHAL* | this study |
| Pet8 | PXMJ19 expressing SSAP from *Corynebacterium variabile DSM 44702* | this study |
| Pet9 | PXMJ19 expressing SSAP from *Escherichia coli MG1655* | this study |
| Pet10 | PXMJ19 expressing SSAP from *Corynebacterium sp HMSC070E08* | this study |
| Pet12 | PXMJ19 expressing SSAP from *Corynebacterium humireducens DSM 45392* | this study |
| Pet13 | PXMJ19 expressing SSAP from *Corynebacterium timonense 5401744* | this study |
