## Supplementary figures and images for "New logic facilitated identification of novel phage recombinase function units"

### S1 Fig

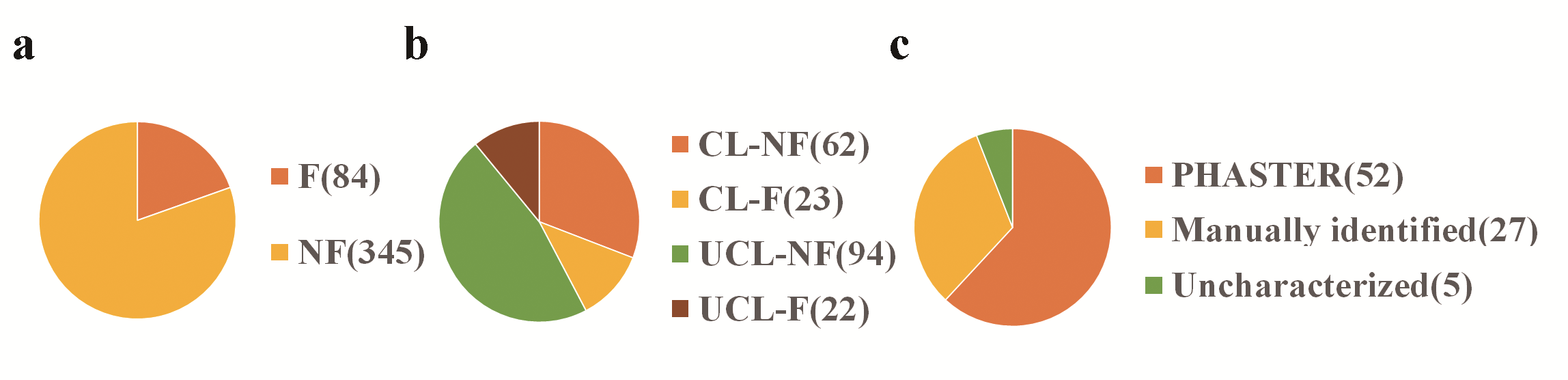

### S2 Fig

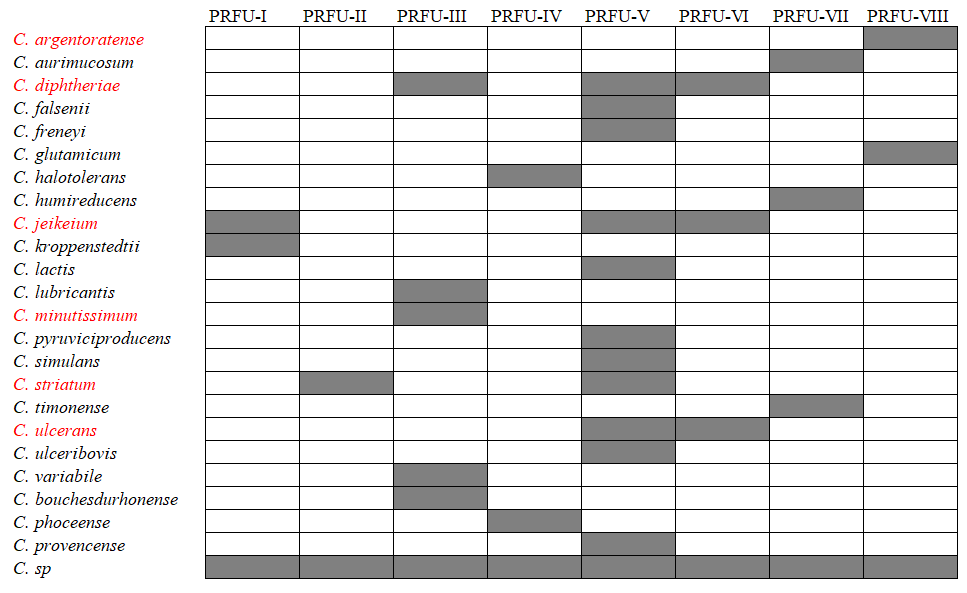

### S3 Fig

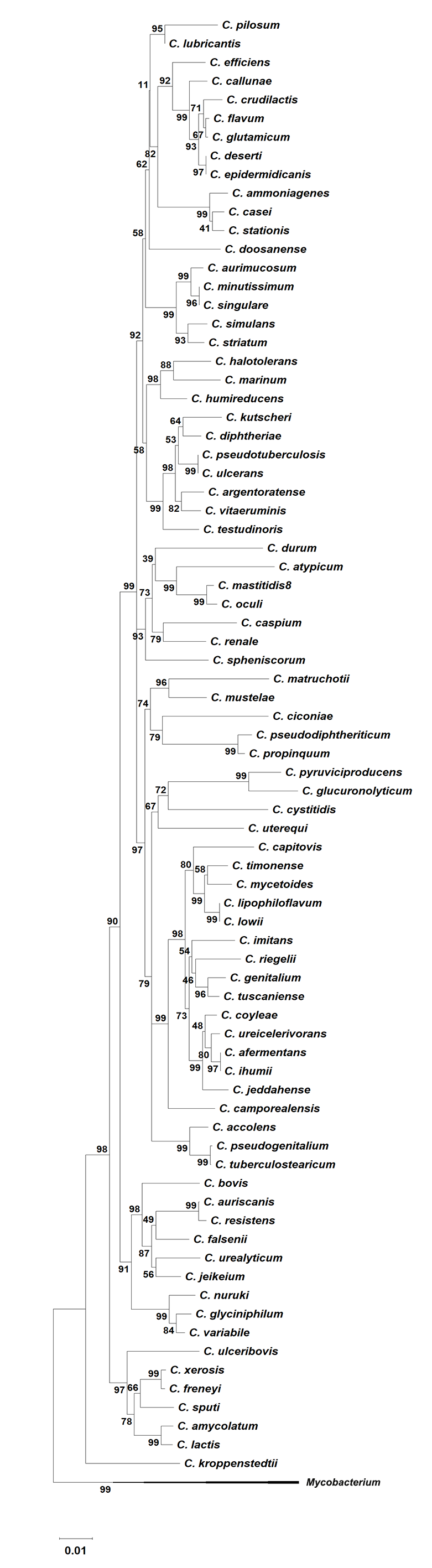

### S4 Fig

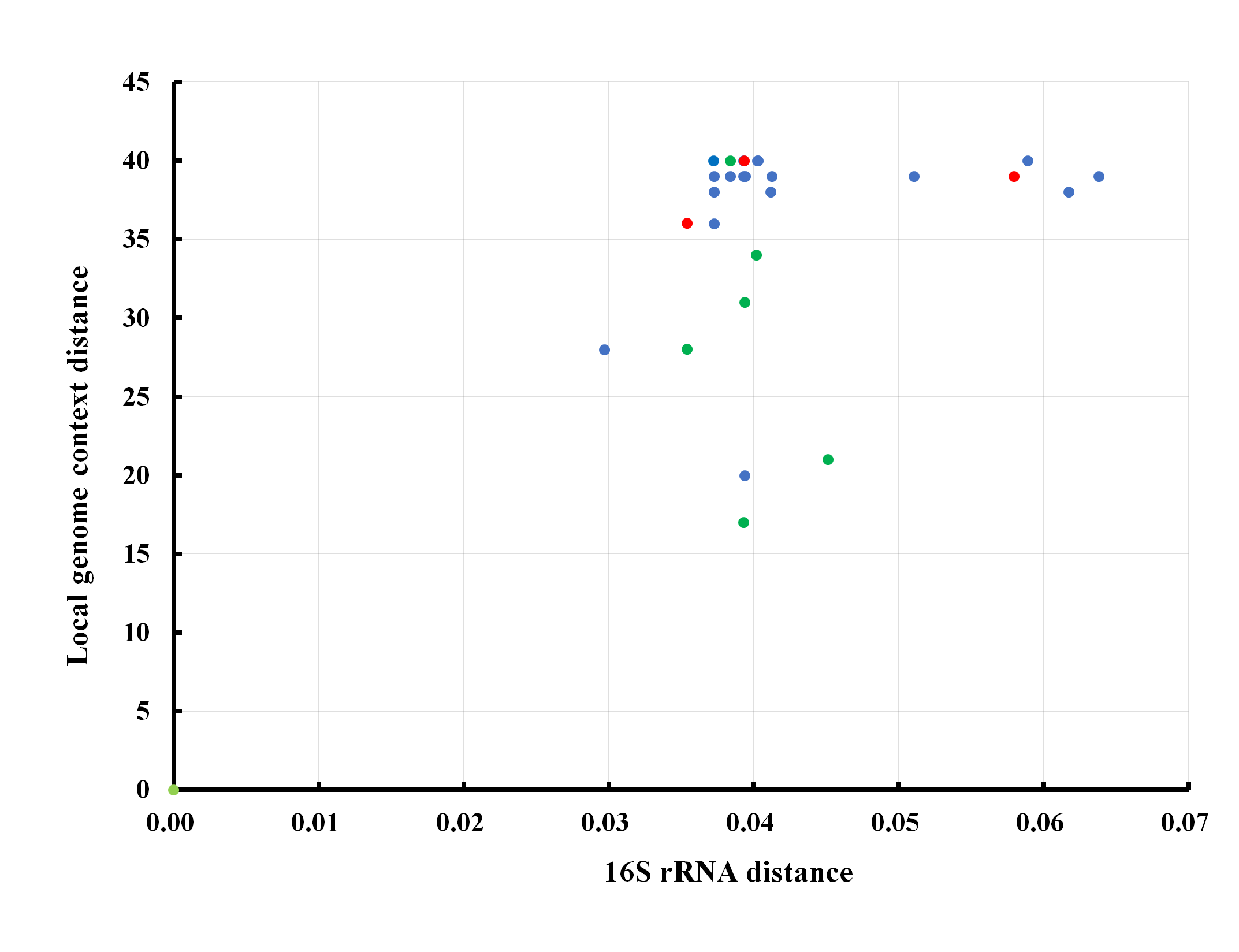
